# Accelerando and crescendo in African penguin display songs

**DOI:** 10.1101/2025.04.04.647185

**Authors:** Taylor A. Hersh, Yannick Jadoul, Marco Gamba, Andrea Ravignani, Livio Favaro

**Author notes:** Joint first authors, corresponding authors. Joint senior authors (Andrea Ravignani, Livio Favaro).

## Abstract

Many species produce rhythmic sound sequences. Some purportedly speed up their vocalizations throughout a display, reminiscent of—but not necessarily equivalent to— “accelerando” in human music. This phenomenon has been frequently reported but rarely quantified, which limits our ability to understand its mechanism, function, and evolution. Here, we use a suite of rhythm analyses to quantify temporal and acoustic features in the display songs of male African penguins (*Spheniscus demersus*). We show that songs get faster (i.e., accelerando) and louder (i.e., crescendo) as they progress. The accelerando occurs because the inter-syllable silences, not the syllables themselves, predictably shorten over time. This rhythmicity is maintained even when individuals take audible breaths. Individuals also show plasticity: when they start with a slow tempo, they speed up more strongly than when they start with a fast tempo. We hypothesize that this well-timed accelerando may stem from arousal-based mechanisms, biomechanical constraints, or more complex rhythmic control; future work should test the mechanisms behind this intra-individual rhythmic variation, since non-passerine birds are thought to have limited vocal plasticity. By integrating a rich empirical dataset with cutting-edge rhythm analyses, we establish the necessary foundation to determine how such features evolved and their role(s) across communication systems.

## 1. Introduction

Theoretically, animals can time their vocalizations in an unlimited number of ways. In practice, isochrony—where the inter-onset intervals (IOIs) of vocalizations exhibit roughly equal durations—dominates across taxa.^1–5^ Temporal variation around isochrony can be achieved by progressively lengthening (ritardando) or shortening (accelerando) IOIs in a sequence. In particular, vocal accelerando has been reported in passerine birds and mammals, and examples are often linked to agonistic interactions and high arousal situations (Table 1). Rufous horneros (*Furnarius rufus*) produce “accelerated trills” when singing solos and duets, with song playing a role in territory defense and mate guarding.^6^ Male thrush nightingales (*Luscinia luscinia*) may use accelerando in mating songs to evoke anticipation in listeners.^7^ Both Columbian and California ground squirrels (*Urocitellus columbianus* and *Otospermophilus beecheyi*, respectively) accelerate call production in the presence of a moving predator or intruder.^8,9^ Yellow-bellied marmots (*Marmota flaviventris*) produce “accelerando whistles” when disturbing objects approach them; these acoustic displays often culminate in a physical attack.^10^ Similarly, when flying male hammer-headed bats (*Hypsignathus monstrosus*) are approached by other males, they accelerate their call rate, orient towards the intruder, and attack if the intruder is not deterred.^11^

**Table 1.** Examples of mammalian (top) and avian (bottom) species with accelerando in vocal displays. Note that most reports of accelerando are anecdotal and descriptive rather than quantitative.

| Animal | Latin name | Vocal display | Reference(s) |
| --- | --- | --- | --- |
| California ground squirrel | *Otospermophilus beecheyi* | Calls | 9 |
| Chimpanzee | *Pan troglodytes* | Pant hoots | 35 |
| Columbian ground squirrel | *Urocitellus columbianus* | Calls | 8 |
| Gibbon | Various species | Great calls | 36,37 |
|  |  | Song | 38 |
| Gorilla | *Gorilla* spp. | Hoot series | 39 |
| Hammer-headed bat | *Hypsignathus monstrosus* | Calls | 11 |
| Rock hyrax | *Procavia capensis* | Song | 15 |
| Yellow-bellied marmot | *Marmota flaviventris* | Accelerando whistles | 10 |
| Australian magpie | *Gymnorhina tibicen* | Song | 40 |
| Australian pied butcherbird | *Cracticus nigrogularis* | Song | 41 |
| Black-chinned sparrow | *Spizella atrogularis* | Song | 42 |
| Common firecrest | *Regulus ignicapilla* | Song | 43 |
| Field sparrow | *Spizella pusilla* | Song | 44,45 |
| Rufous hornero | *Furnarius rufus* | Accelerated trills in song | 6 |
| Song sparrow | *Melospiza* spp. | Trills (especially introductory trills) | 46 |
| Thrush nightingale | *Luscinia luscinia* | Song | 7,47 |
| Western capercaillie | *Tetrao urogallus* | Calls | 48 |
| Wilson's snipe | *Gallinago delicata* | Song | 43 |
| Wood warbler | *Phylloscopus sibilatrix* | Song | 42 |
| Wrentit | *Chamaea fasciata* | Song | 42 |

More broadly, there is evidence that loud, accelerating calls are an ancestral call morphotype for phylogenetically new and ancient primates.^12,13^ Accelerando is also common in human music, where it is used as a stylistic device to increase arousal, manipulate tension, and prevent habituation in listeners.^14–16^ Coupling accelerando with other musical phenomena, such as crescendo (i.e., increasing loudness/intensity), can enhance a composition’s expressiveness.

These and other descriptions of accelerating animal acoustic displays suggest that accelerando plays an important role in diverse communication systems, although that role may not necessarily be equivalent across systems. However, questions relating to the evolution, mechanism, and function of accelerando will remain intractable unless one can explicitly and systematically quantify accelerando across diverse study systems.^17^ Here, we use a suite of rhythm analyses and statistical models to measure various temporal and acoustic features, including accelerando, in vocalizations produced by a non-passerine bird, the critically endangered African penguin (*Spheniscus demersus*). Compared to oscine songbirds, vocal communication is understudied in other avian taxa, especially in those assumed to have limited vocal plasticity.^18^

African penguins are territorial, monogamous seabirds. They live in dense colonies where they compete for nesting space. During breeding periods, individuals produce multisyllabic “ecstatic display songs” (hereafter *songs*; Figure 1).^19,20^ Primarily made by males, these songs seem to play a role in nest territory defense, mate recognition, and mate attraction, but how they bolster or facilitate such displays is unknown.^19–22^ Songs are typically composed of a sequence of short-duration “A” syllables leading up to one or more long-duration “B” syllables.^20^ Both A and B syllables are egressive (i.e., produced with an outward flow of air). A third syllable type, the ingressive “C” syllable, occurs when penguins audibly inhale. C syllables can be interspersed throughout a song but do not occur in all songs. Given the differences in how and when they are produced, whether C syllables contribute to the structure and function of songs in the same way as A and B syllables is unknown.

**Figure 1.**
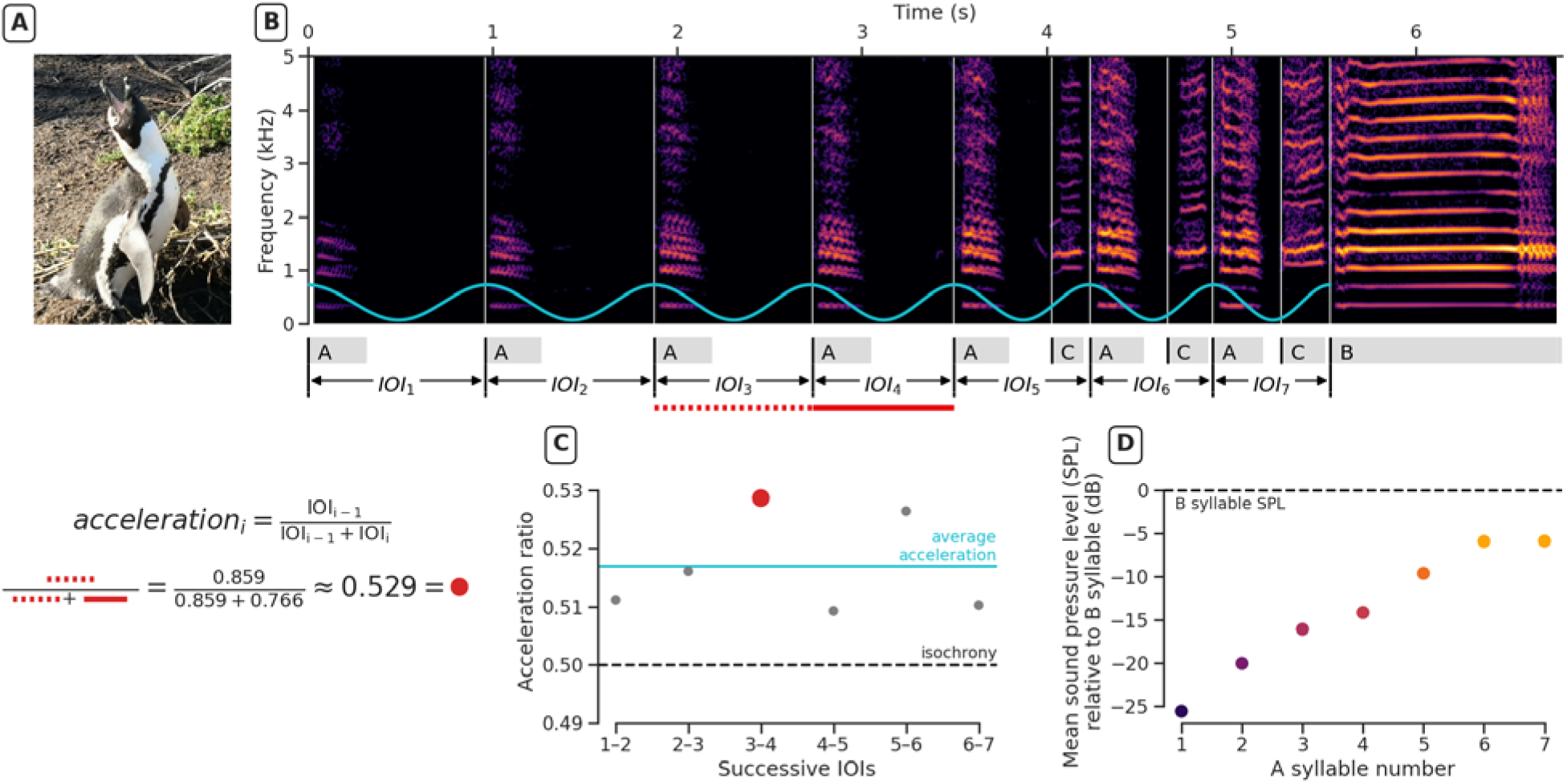
Example of accelerando and crescendo in an African penguin ecstatic display song. **(A)** Typical penguin stance while singing, with feet apart, beak up, and wings extended. **(B)** The annotated spectrogram shows the portion of a song leading up to the first B syllable (Audio S1). The accelerating sine wave (blue) captures the acceleration of the A syllables. As this song progresses, **(C)** successive IOIs shorten (accelerando) and **(D)** successive syllables get louder (crescendo).

Temporal features of vocalizations often have higher transmission fidelity than spectral features in noisy environments, such as dense colonies of highly vocal individuals.^23^ This could evolutionarily favor information encoding in the time domain for species like penguins and motivated our analysis of the temporal structure of African penguin songs. While spectral plasticity in African penguin vocalizations is limited, temporal plasticity is virtually unexplored.^21^ Understanding song structure is thus a critical first step towards eventually unravelling the exact function of this biologically important display.

Using song recordings from ex-situ individuals, we first test whether C syllables contribute to the temporal structure of songs in the same way as A syllables. Then, we investigate whether qualitative impressions of song accelerando and crescendo are quantitatively validated. We interpret our results against the broader behavioral ecology of African penguins and conclude with topics for future research.

## 2. Materials and Methods

### (a) Song recordings

We analyzed 551 passively recorded songs from 26 ex-situ male African penguins (Table S1) during the 2016/2017 breeding seasons (between October 2016 and May 2017) at three Italian zoological facilities. Each male contributed at least six songs. Recordings were made when the facilities were closed to visitors, thus minimizing activity and disturbance around the exhibits. Individuals were unequivocally identified using colored flipper bands. Our dataset is a subset of a previously collected one, excluding females; data collection methods are detailed in ^22^ and summarized in Method S1.

### (b) Song feature extraction

Song spectrograms were visually inspected and manually annotated in Praat^24^ version 6.0.43 by two trained analysts. Each syllable’s start and end were manually demarcated and syllables were assigned to a syllable type based on spectrotemporal features, as previously described in the literature.^19,20,25^ Inter-observer agreement on syllable type assignments was high (Cohen’s κ coefficient = 0.99; see ^22^ for details). Start time and duration were automatically extracted from the annotations with a custom Python script for each of the *n* syllables up to and including the first B syllable in a song. Additionally, each syllable’s intensity was calculated with the Python package Parselmouth^26^ version 0.4.1 (Praat version 6.1.38) as the energy-based average of the intensity curve using default parameters after filtering out low-frequency noise using a Hann band-stop filter below 100 Hz (bandwidth 10 Hz).

We then calculated the *n* −1 IOIs and silent interval durations between successive syllables. For each song, we calculated acceleration ratios between successive IOIs as ^4^

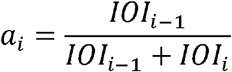

(resulting in *n* −2 ratios, Figure 1) and average acceleration ratio (excluding □ □ □_1_) as:

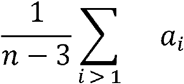

A ratio greater than 0.5 corresponds to an accelerating sequence (i.e., a sequence with shortening IOIs and thus an increasing tempo), a ratio equal to 0.5 corresponds to an isochronous sequence, and a ratio less than 0.5 corresponds to a decelerating sequence.

To normalize the position of syllables (or IOIs, or acceleration ratios) over songs of different lengths, we calculated their relative position within a song such that it is scaled between 0% and 100% for the first and last syllable (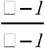 for *syllable*_*i*_, 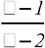 for *IOI*_*i*_, 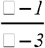 for *a*_*i*_).

Finally, for each penguin, we grouped the IOI durations into five equally spaced bins based on their relative position (see Method S2). We then calculated the mean, standard deviation, and coefficient of variation (CV, corrected for small sample size)^27^ of the IOI durations of all songs within each bin.

### (c) C syllable analysis

Our first goal was to determine whether A and C syllables contribute equally to the acceleration of a song. This analysis assessed whether we should include C syllable onsets in IOI calculations, acceleration ratio calculations, and subsequent analyses. To address this, we calculated acceleration ratios for all pairs of successive IOIs and generated three distributions for comparison. The baseline distribution includes only acceleration ratios in parts of songs *without* C syllables present (i.e., between successive A-A and A-B syllable IOIs). The two alternative distributions include acceleration ratios in parts of songs *with* C syllables present. For these two alternative distributions, the C syllable onsets were either omitted or included in the IOI and acceleration calculations. We call these the “C-omitted distribution” and the “C-included distribution”, respectively. We then compared the alternative distributions to the baseline distribution to see which one was more similar to the baseline distribution. To do so, we calculated a kernel density estimation of the acceleration ratios in the baseline distribution and used it to calculate the likelihood of all acceleration ratios in the C-omitted and C-included distributions. We then used a Wilcoxon signed-rank test to test the difference in the median likelihood of each penguin’s acceleration ratios, omitting vs. including C syllable onsets. These tests were only done for penguins with at least three C syllables in their songs (*n*=13). The baseline distribution represents the empirical distribution of measured acceleration ratios in the absence of detectable C syllables; by comparing which alternative distribution matches the baseline distribution most closely, we can quantitatively assess which of the two approaches, omitting or including C syllable onsets, can explain the observed temporal patterns in the most parsimonious way.

### (d) Song accelerando and crescendo analyses

To determine whether songs accelerate, we ran a linear mixed-effects model (LMEM) (response variable: log-transformed IOI duration; fixed effect: relative position of IOI within song; nested random intercept: song in penguin in colony). As a linear slope in log space corresponds to an exponential curvilinear relation in the non-transformed data, the coefficient fitted by the LMEM conceptually matches the acceleration ratio between successive IOIs. For this and other LMEMs, we tested the significance of the fixed effect using a chi-square likelihood ratio test, comparing model fits with vs. without the fixed effect.

Having quantitatively shown accelerando, we next ran a LMEM to determine whether acceleration is constant throughout a song (response variable: acceleration ratio; fixed effect: relative position of acceleration ratio within song; nested random intercept: song in penguin in colony). We also wanted to know how the two components of an IOI—syllable and silence— contribute to the observed acceleration. Accordingly, we ran LMEMs analogous to the previous ones, checking whether the duration of syllables and silences changes throughout songs (response variable: log-transformed syllable/silence duration; fixed effect: relative IOI position; nested random intercept: song in penguin in colony).

To determine whether IOI durations become more precise as songs progress, we fitted a LMEM to test the change in IOI duration CV across songs (response variable: CV of grouped IOI durations; fixed effect: relative position of IOI bin; nested random intercept: penguin in colony). CV values were calculated for 5 disjunct bins; this approach groups values to calculate the CV summary statistic but also quantifies its evolution across songs. We also tested the decrease in CV between IOI_1_ (the first IOI) and IOI_n-1_ (the last IOI before the B syllable) for each individual penguin using the modified signed-likelihood ratio test (SLRT) for equality of CVs, implemented in the R package *cvequality*.^28,29^

Next, we ran a LMEM to test whether IOI_1_ predicts average acceleration in songs (response variable: average acceleration ratio, excluding IOI_1_; fixed effect: IOI_1_ duration; nested random intercept: penguin in colony). As our random effect explicitly controls for variation across individuals (i.e., between penguins), the tested fixed effect captures within-penguin variation.

Finally, we wanted to know whether songs crescendo leading up to the first B syllable. Again, we tested this with a LMEM (response variable: A syllable intensity, normalized relative to the first B syllable intensity; fixed effect: relative syllable position in the song; nested random slope: song in penguin in colony).

## 3. Results

When C syllable onsets are included in IOI calculations, the distribution of acceleration ratios between successive IOIs (C-included distribution) clearly differs from the observed acceleration in naturally occurring sequences of A/B syllables without any Cs (baseline distribution; Figure 2A). When C syllable onsets are omitted (i.e., not factored into IOI and acceleration ratio calculations), the resulting acceleration ratios (C-omitted distribution) match the baseline distribution significantly better. More specifically, the likelihood of the C-omitted distribution data points being drawn from the baseline distribution is significantly higher than for the points from the C-included distribution (Wilcoxon signed-rank test, *n*=13 penguins, *T*=0.0, *p*<0.001; Figures 2A & S1). These results indicate that the underlying assumption of the C-omitted distribution—that C syllables do not fulfil the same role as A syllables in a song’s temporal structure—is the more parsimonious explanation for the observed accelerando. Accordingly, we omitted C syllables in subsequent analyses.

**Figure 2.**
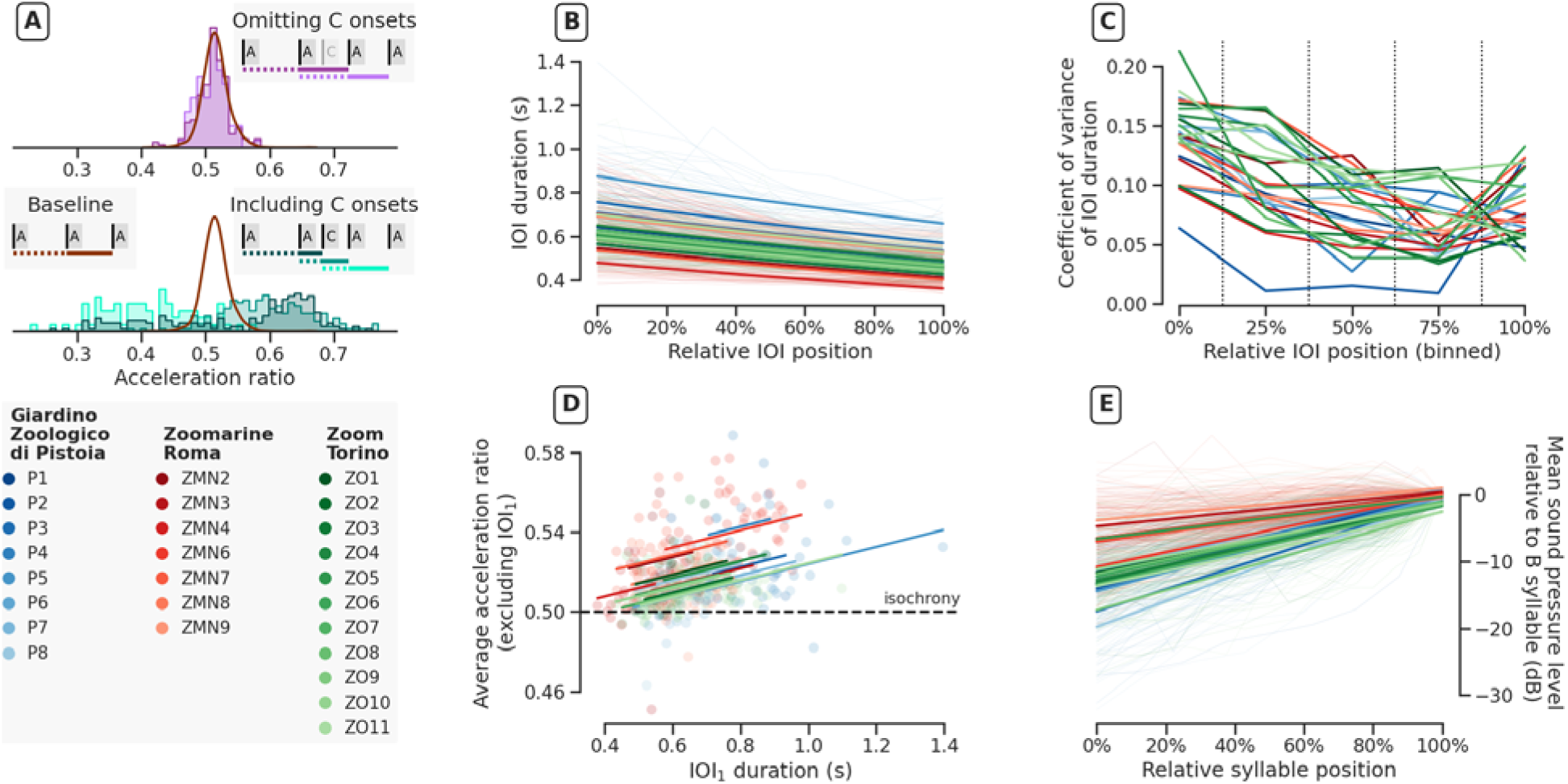
Quantitative analyses of accelerando and crescendo in African penguin ecstatic display songs. **(A)** Omitting C syllable onsets results in a distribution of acceleration ratios (purple) closely matching the baseline distribution (brown curve) of acceleration ratios between two successiv egressive (A/B) syllable IOIs (top); including C syllable onsets (teal) does not (bottom). This suggests that C syllables do not contribute to the temporal structure of songs in the same way a A syllables. **(B)** Within a song, IOI duration is significantly predicted by its relative position. The background lines show the individual songs; the bold lines show the fitted LMEM’s effect for each penguin (excluding the nested random effect per song). **(C)** Across penguins, the coefficient of variation of IOI duration significantly decreases as songs progress. The vertical lines represent the boundaries between bins (see Method S2). **(D)** Controlling for individuals, a LMEM show that IOI_1_ significantly predicts a song’s subsequent acceleration. **(E)** Songs also show crescendo: later syllables are louder than earlier syllables. Bold lines again show the LMEM fit for each penguin. In panels B–E, colors denote different penguins (see bottom left legend; alphanumeric codes from ^22^) and shades correspond to the three colonies.

Next, our analyses show that African penguin songs exhibit robust accelerando. Across penguins, an IOI’s position in a song is significantly negatively correlated with its duration, such that later IOIs have shorter durations (chi-square test, *n*=2834 syllables, χ^*2*^(1)=2425.2, *p*<0.001; Figure 2B). Per-penguin linear regressions confirm these results: IOIs shorten as songs progress for all 26 penguins (Figure S2).

Song acceleration is above 0.5 but not constant, with smaller acceleration ratios at the end of songs compared to the start (chi-square test, *n*=2253 syllables, χ^*2*^(1)=65.861, *p*<0.001; Figure S3). In general, penguins do not sing one stereotyped song, instead showing within-individual plasticity in the number of A syllables, number of C syllables, and mean acceleration ratio per song (Figure S4).

Our duration analyses show that shortening silences, not A syllables, cause the observed acceleration. IOI position is negatively correlated with silence duration, with IOIs becoming ∼25% shorter and inter-syllable silences decreasing by almost 45% as songs progress (chi-square test, *n*=2834 syllables, χ^*2*^(1)=3211.3, *p*<0.001; Figure S5). In contrast, IOI position is positively correlated with A syllable duration: A syllables that occur later in songs are ∼20% longer than A syllables that occur earlier in songs (chi-square test, *n*=2834 syllables, χ^*2*^(1)=632.55, *p*<0.001; Figure S5).

Across songs, the IOI durations become significantly less variable throughout songs (chi-square test, *n*=130 CV measurements, χ^*2*^(1)=56.739, *p*<0.001; Figure 2C). In other words, songs gradually get more precise towards the end than at the start. Per individual, 13/26 penguins show a significant decrease in CV when comparing IOI_1_ to IOI_n-1_ (Figure S6).

Moreover, the duration of IOI_1_ predicts the average acceleration ratio leading up to the first B syllable: a longer IOI_1_ is correlated with a larger acceleration (Figure 2D; chi-square test, *n*=465 songs, χ^*2*^*(1)*=47.102, *p*<0.001). Again, per-penguin linear regressions broadly confirm these results (Figure S7). Figure S8 illustrates this correlation for one penguin, showing the acceleration for ten songs that vary in the length of IOI_1_.

Finally, the intensity of A syllables significantly increases with relative syllable position, showing that songs crescendo leading up to the first B syllable, which is typically the most intense syllable in a song (chi-square test, *n*=3357 A and B syllables, χ ^*2*^*(1)*=7.571, *p*<0.01; Figure 2E). Per-penguin linear regressions confirm this result, with significant positive slopes for all 26 penguins (Figure S9).

## 4. Discussion

The animal communication literature contains many descriptions of accelerating vocal signals (Table 1), but few studies explicitly quantify accelerando. Our analyses show that male African penguin songs get faster and louder as they progress. They are also more precise (regarding IOI duration) at the end compared to at the start. The accelerando occurs because males shorten the silences between A syllables as they sing and is made more impressive by the fact that the A syllables themselves become longer throughout a song. While the accelerando itself is not perfectly constant, songs consistently accelerate rather than become isochronous or slow down. The crescendo occurs because males produce increasingly intense A syllables, culminating in a very long and intense B syllable.

Past work shows that C syllables differ from A syllables in how they are produced and how consistently they occur in song.^20^ Our results extend that work, suggesting that C syllables do not contribute to song accelerando in the same way as A syllables. Whether these two syllable types are also functionally different is an open question; playback experiments that evaluate receiver response to songs where A and C syllables are added or removed are needed to conclusively test this. Regardless, our results suggest that C syllables should be omitted from African penguin song analyses specifically focused on temporal structure.

The duration of the first IOI predicts subsequent acceleration in a song. In other words, natural variation in IOI_1_ is accounted for by a correspondingly more or less strong acceleration ratio throughout a song. This is not explained by individual penguins filling specific acceleration “niches” (for example, one penguin always accelerates slowly while another penguin always accelerates quickly), as evidenced by penguins overlapping in their song parameter distributions. Coupled with the increasing precision of IOIs as the song progresses, these results suggest that individual penguins show some degree of vocal plasticity in the temporal domain.

Which mechanisms can explain the predictable accelerando and crescendo we observe in African penguin songs? Arousal is one possibility, but the proximal mechanisms linking arousal to changes in call speed or intensity are often difficult to determine.^30^ They can include: increased muscle tone that alters the resonant properties of the underlying vocal production anatomy; an increased neural duty signal through the vocal motor regions in the brain; or an increase in the rate of other linked physiological processes, such as heart rate.^31–33^ From a biomechanical perspective, the patterns we observe match what we would expect if penguins are physically preparing to produce the intense, long B syllable when singing. For example, if a certain air pressure level or volume in the lungs must be reached before a B syllable can be produced, then penguins may incrementally work towards that threshold, gradually adding air to their lungs and increasing tension in their syrinx membranes leading up to producing the B syllable. This scenario could explain the patterns we see in C syllable production and A syllable crescendo. Alternatively, African penguins may aim for a rhythmic “target” when singing, e.g., a specific tempo, and compensate for any natural variation in IOI_1_ by accelerating more or less strongly. Under this scenario, our finding that C syllables do not contribute to temporal song structure in the same way as A syllables could indicate that penguins time their breaths in such a way as not to delay the onset of the next A syllable and thus interrupt the rhythmicity/acceleration of their song. This could be analogous to human wind instrument musicians, who are taught to breathe during rests (i.e., non-note moments) in a musical performance. If penguins instead breathed randomly without compensating for the breath duration, some of their breaths would overlap with or A delay syllables, altering the temporal structure of the vocal display.

To tease apart these alternative hypotheses, future work should collect physiological and anatomical data from singers in parallel with song measurements to look for correlations between singer body condition/quality and spectrotemporal song features. This would enable testing whether arousal is responsible for the observed accelerando and crescendo in African penguin songs. Playback experiments that manipulate different song features and measure conspecific responses will shed light on which features are salient and which individuals are the intended signal receiver(s). Coupling such experiments with real-time physiological monitoring (e.g., heart rate)^34^ of nearby penguins will help determine whether song accelerando and crescendo influence arousal, tension, and/or habituation in listeners, as they do in human music.^14,16^

Our work adds African penguins to a growing list of animal species that produce accelerating vocal displays and takes a step forward by explicitly quantifying, rather than describing, this acoustic phenomenon. Our approach can help researchers conduct similar quantitative analyses for other species’ vocal displays. By systematically quantifying accelerando—a fundamental feature of rhythmic communication—and crescendo in animal vocalizations, we can establish the necessary foundation to determine how such features evolved and understand their role(s) across communication systems.

## Supporting information

Full supplement

## Ethics

Under Italian law, no specific permissions were required for collecting non-invasive passive acoustic recordings of ex-situ penguins.

## Data accessibility

The dataset supporting the conclusions of this manuscript is available in a dedicated Open Science Framework repository (https://osf.io/4dfxz/?view_only=979c3a6ae90942beb74269856b9504de).

## Authors’ contributions

All authors were involved in conceptualization; TAH, YJ, and AR developed the methodology; YJ created the Python scripts for the acoustic and statistical analyses; TAH and YJ analyzed the data and prepared the visualizations; MG, AR, and LF supervised the research; LF provided project resources; AR and LF acquired funding; TAH and YJ wrote the original draft; all authors reviewed and edited the final draft.

## Acknowledgements

We thank Giardino Zoologico di Pistoia, Zoomarine Roma, and Zoom Torino for facilitating research and providing access to penguin colonies; Stephanie King, Elias Fernández Domingos, Christian Herbst, Scott McCain, and two reviewers for helpful feedback; Eleonora Cresta and Elena Fumagalli for annotating recordings; and Francesca Terranova for checking text grids.

