## Supplementary material for "Accelerando and crescendo in African penguin display songs": Full supplement

**Accelerando and crescendo in African display penguin songs – Supplementary material**

*Method S1. Song recording protocol*

Recordings were made in outdoor exhibits that ranged in size from 70–1500 m^2^. Penguins were recorded using a RØDE NTG-2 super-cardioid microphone (frequency response: 20 Hz–20 kHz, sensitivity: −36±2 dB re 1 V/Pa at 1 kHz, maximum sound pressure level: 131 dB) on a RØDE PG2 pistol grip. The microphone and grip were placed 5–10 m from vocalizing individuals. The microphone output signal was digitized using a TASCAM portable recorder (model: DR-680 or DR-40, sampling rate: 44.1 kHz) and saved to an internal SD memory card in WAV format (resolution: 16 bits). Songs from female penguins are rare^1,2^ and were excluded because the sample size was too small to statistically account for any potential sex effect in our analyses. The number of penguins in each colony varied throughout our study (due to deaths, hatches, etc.), but was approximately 23 for Giardino Zoologico di Pistoia, 14 for Zoomarine Roma, and 66 for Zoom Torino. Note that captive African penguins generally have a more flexible breeding cycle than their wild counterparts.^3,4^

*Method S2: Relative IOI position binning protocol*

When calculating how the coefficient of variation (CV) of IOI duration changes throughout songs, the different lengths of the songs complicate grouping the relative positions together. For example, a song with three IOIs will have IOI durations at relative positions 0%, 50%, and 100%, whereas a song with four IOIs will have IOI durations at relative positions 0%, 33%, 66%, and 100%. To be able to calculate how the CV changes throughout songs, we grouped the IOIs into five bins, around 0%, 25%, 50%, 75%, and 100%, and assigned each IOI duration to its respective bin. More precisely, this results in the following bins: [0, 0.125); [0.125, 0.375); [0.375, 0.625); [0.625, 0.875); [0.875, 1].

*Audio S1 (separate files). Audio file and annotations of the ecstatic display song featured in Figure 1*

This ecstatic display song was produced by penguin P5 from the Giardino Zoologico di Pistoia colony (Table S1). The full audio file (~12.5 s) and corresponding Praat text grid with syllable annotations is provided, but only the portion leading up to and including the first B syllable (~7 s) was included in our analyses.

*Table S1. Number of ecstatic display songs recorded from captive male African penguins included in our study*

Penguin identity codes from ^2^ and from the European Association for Zoos and Aquariums (EAZA) are listed for each male.

| *Colony* | *Code from* ^2^ | *Code from EAZA* | *Number of songs* |
| --- | --- | --- | --- |
| Giardino Zoologico di Pistoia | P1 | 10409 | 8 |
|  | P2 | 6100 | 8 |
|  | P3 | 2207 | 10 |
|  | P4 | 2592 | 9 |
|  | P5 | 8199 | 26 |
|  | P6 | 9173 | 24 |
|  | P7 | 7215 | 27 |
|  | P8 | 14140 | 24 |
| Zoomarine Roma | ZMN2 | 9060 | 14 |
|  | ZMN3 | 10092 | 23 |
|  | ZMN4 | 13255 | 21 |
|  | ZMN6 | 13055 | 46 |
|  | ZMN7 | 3852 | 33 |
|  | ZMN8 | 13252 | 42 |
|  | ZMN9 | 9061 | 52 |
| Zoom Torino | ZO1 | 4109 | 6 |
|  | ZO2 | 13124 | 8 |
|  | ZO3 | 3930 | 9 |
|  | ZO4 | 3867 | 7 |
|  | ZO5 | 7003 | 7 |
|  | ZO6 | 770 | 10 |
|  | ZO7 | 4176 | 13 |
|  | ZO8 | 7006 | 15 |
|  | ZO9 | 3866 | 11 |
|  | ZO10 | 773 | 29 |
|  | ZO11 | 4110 | 69 |
| **Total** | **26 males** | | **551 songs** |

*Figure S1. Analysis and test of C syllables’ contribution to accelerando*

When the C syllable onsets are omitted from acceleration ratio calculations, the ratios have a higher likelihood to be drawn from the baseline distribution compared to when the C syllable onsets are included. The median log-likelihood of both distributions, C-omitted and C-included, is significantly different (Wilcoxon signed-rank test, *n*=13 penguins, T=0.0, *p*<0.001; see main text and Figure 2A), indicating that C syllables do not fulfil the same role as A syllables in a song’s temporal structure. Colors denote penguin identity (Figure 2).

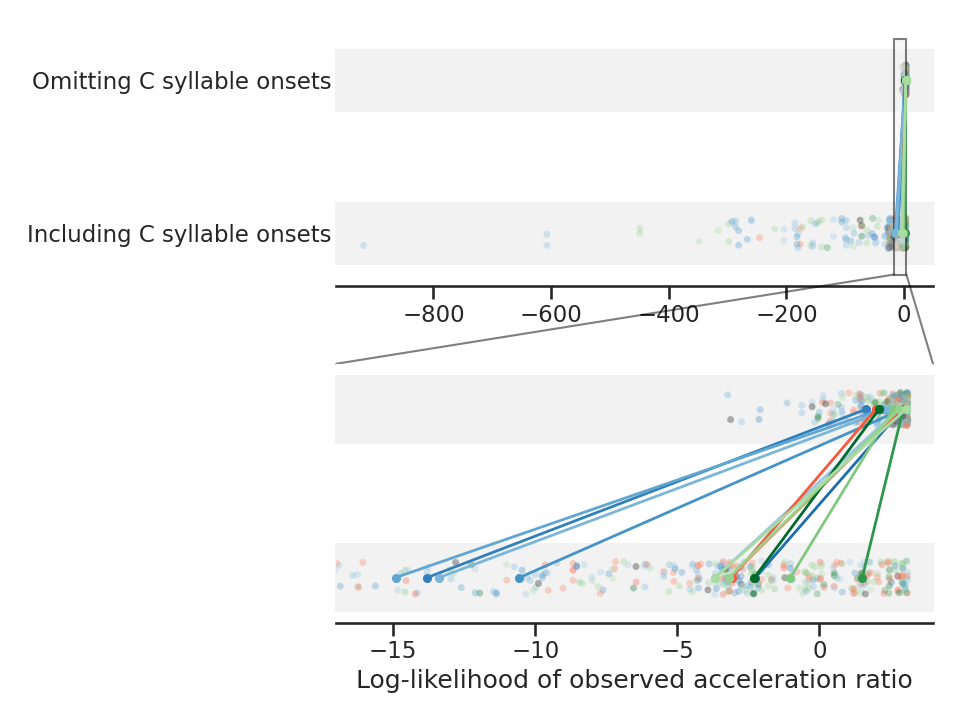

*Figure S2. Per-penguin linear regressions for relative IOI position vs. log-IOI*

For each of the 26 penguins, the fitted linear regression using all song syllables is highly significant. This confirms the result we find with a single LMEM (see main manuscript): the IOI in a penguin’s song is negatively correlated with its relative position within a song. Colors (Figure 2) and alphanumeric codes (Table S1) denote penguin identity.

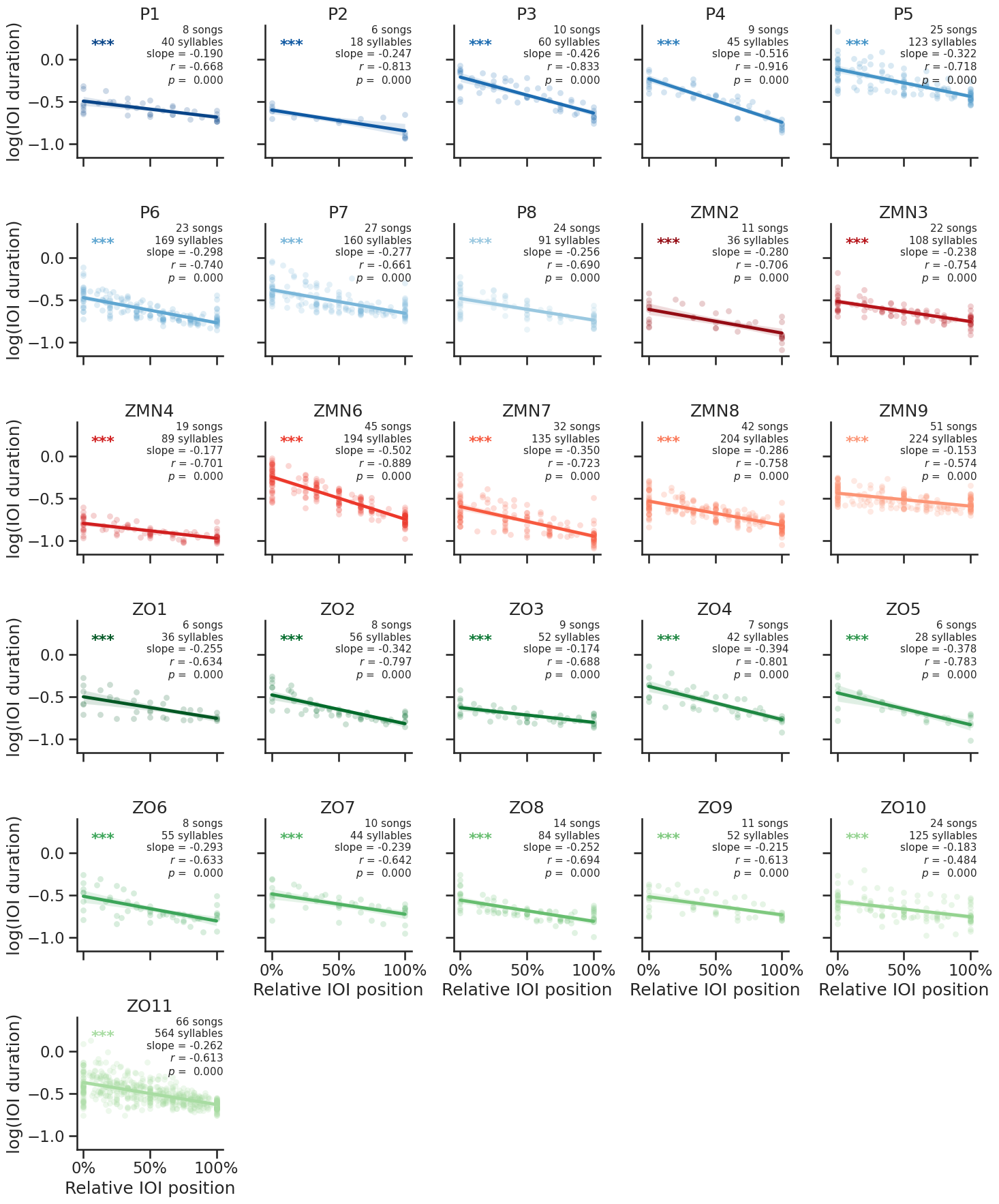

*Figure S3. Analysis and test of the change in acceleration across songs*

Across penguins, songs accelerate from the start to the end, though acceleration is not completely constant. **(A)** Acceleration is significantly lower at the at the end of songs than at the start (chi-square test, *n*=2253 syllables, *χ^2^*(1)=65.861, *p*<0.001), but the effect is small (LMEM estimate: -0.00831 from start to end). Colors denote penguin identity (Figure 2). **(B)** After splitting the acceleration ratios into five bins (see Method S2 for details) based on their relative position within the song, the resulting distribution of acceleration ratios confirms this. In all five bins, the acceleration ratios are significantly higher than 0.5 (i.e., the ratio corresponding to isochrony), showing consistent accelerando (five Wilcoxon rank sum tests; *p* < 0.001 for all). Per each bin, the fraction of acceleration ratios > 0.5, from left to right, is: 83.8%, 79.9%, 82.2%, 81.4%, and 69.6%.

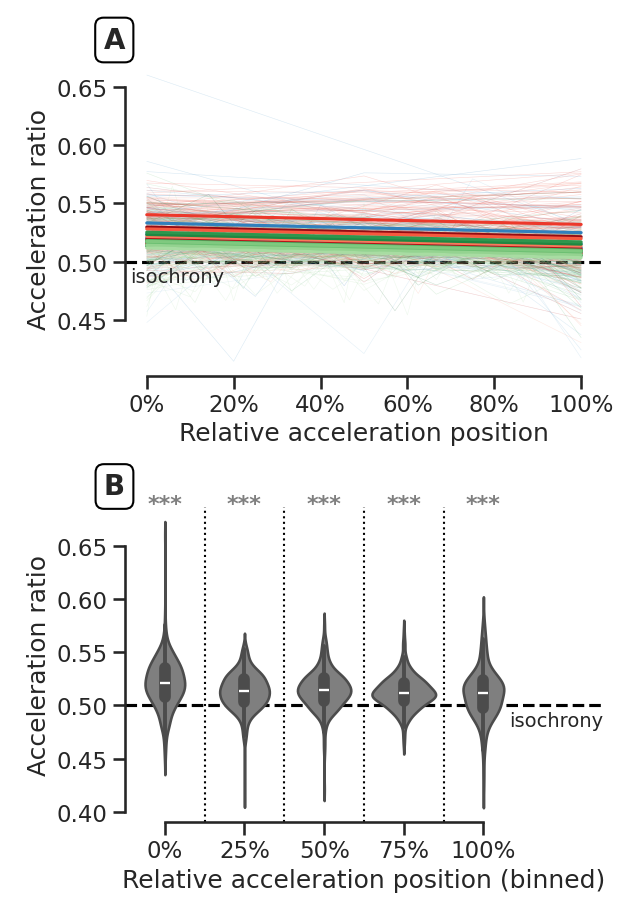

*Figure S4. Between- and within-individual variation in song structure and mean acceleration*

Several song characteristics vary strongly within penguins, indicating that each penguin does not have a fixed song structure. This is shown by the per-penguin distributions of **(A)** the number of A syllables in a song, **(B)** the number of C syllables in a song, and **(C)** the mean IOI acceleration ratio per song. Not all penguins have an identical range of values, but their distributions overlap considerably and demonstrate that penguins show a certain level of vocal plasticity. Colors (Figure 2) and alphanumeric codes (Table S1) denote penguin identity.

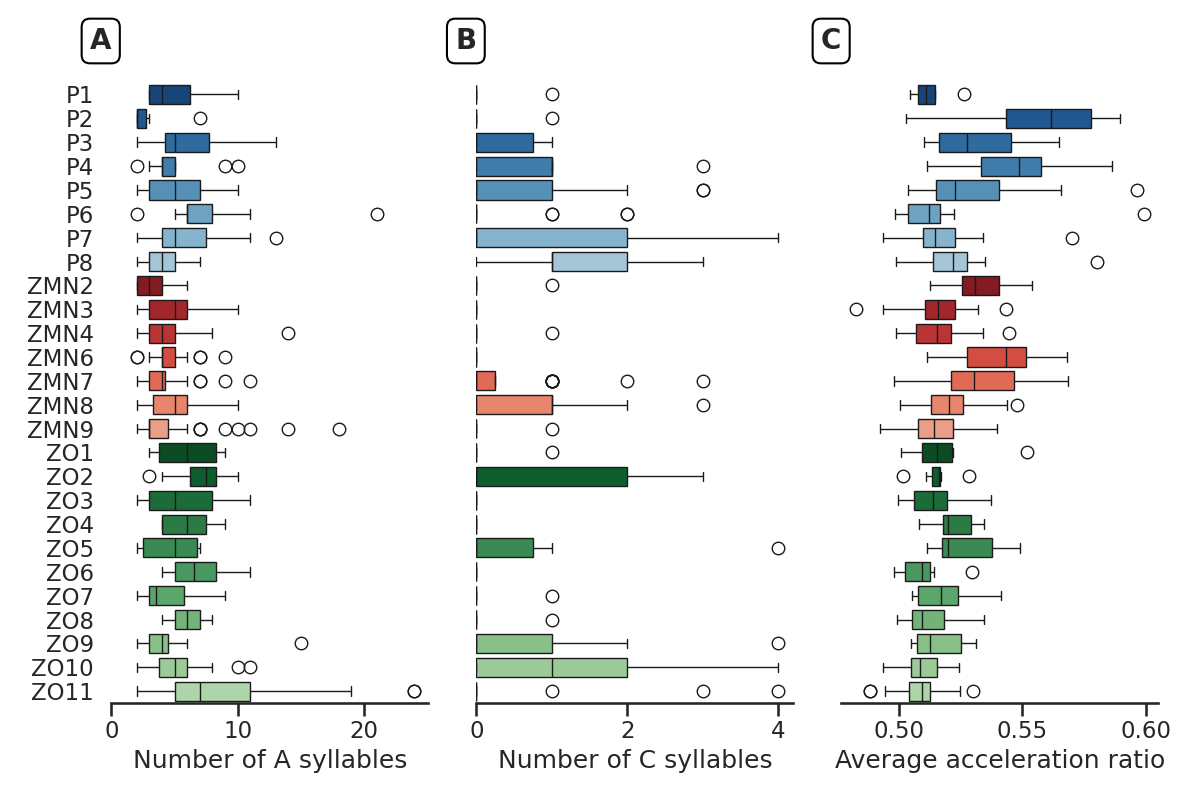

*Figure S5. Analysis and tests of the change in duration of IOI components (silences and syllables) as songs progress*

LMEMs show the duration of inter-syllable silences decreases throughout songs, whereas the duration of A syllables increases. **(A)** Relative IOI position is negatively correlated with silence duration, with IOIs becoming ~25% shorter and inter-syllable silences decreasing by almost 45% as songs progress (chi-square test, *n*=2834 syllables, *χ^2^*(1)=3211.3, *p*<0.001). **(B)** In contrast, relative IOI position is positively correlated with A syllable duration: A syllables that occur later in songs are ~20% longer than A syllables that occur earlier in songs (chi-square test, *n*=2834 syllables, *χ^2^*(1)=632.55, *p*<0.001). Colors denote penguin identity (Figure 2).

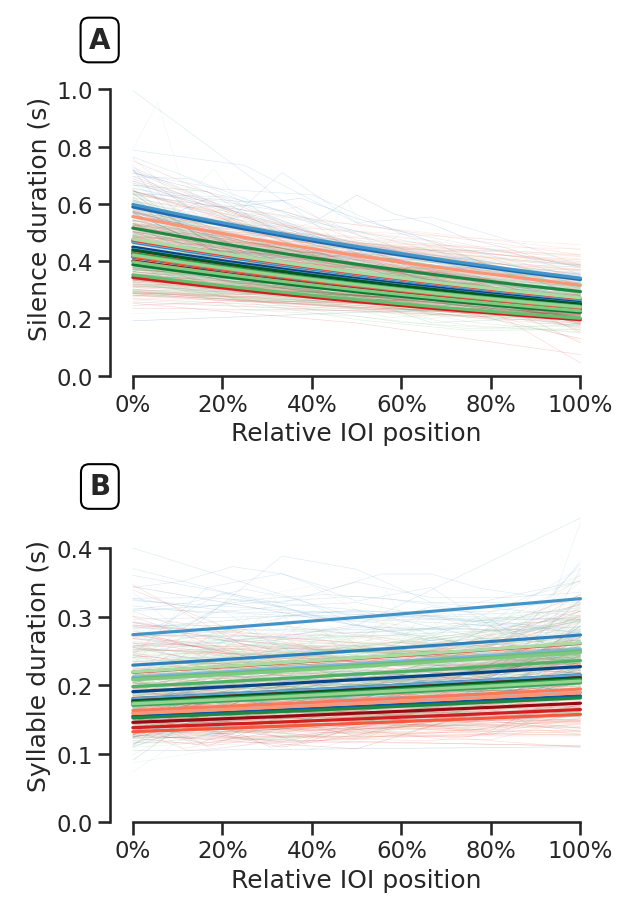

*Figure S6. Per-penguin boxplots of the distribution of IOI_1_ and IOI_n-1_*

Boxplots of each penguin’s IOI_1_ and IOI_n-1_ illustrate the reduced variance in IOI duration between the start and the end of a song. Using the modified signed-likelihood ratio test (MSLRT) to compare coefficients of variation (CVs), 13 out of 26 penguins show a significant decrease in CV, even given the limited number of songs for some penguins. Colors (Figure 2) and alphanumeric codes (Table S1) denote penguin identity.

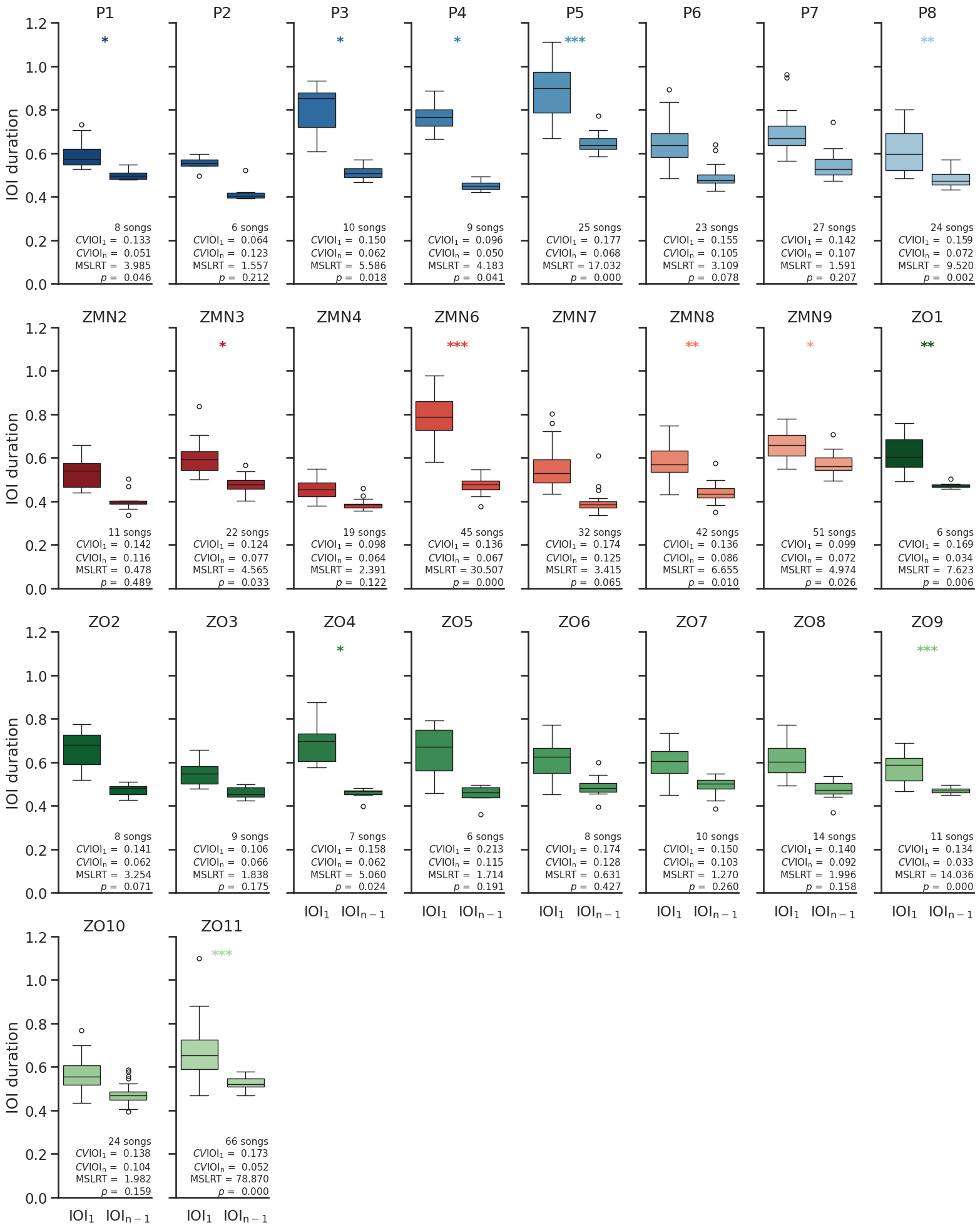

*Figure S7. Per-penguin linear regressions for IOI_1_ vs. average acceleration after IOI_1_*

Seven out of 26 fitted linear regressions—one per penguin—show a significant positive correlation between IOI_1_ and the average acceleration after IOI_1_; 22/26 have a positive slope. This supports the significant result we find when fitting a single LMEM on the combined data (see main manuscript). Colors (Figure 2) and alphanumeric codes (Table S1) denote penguin identity.

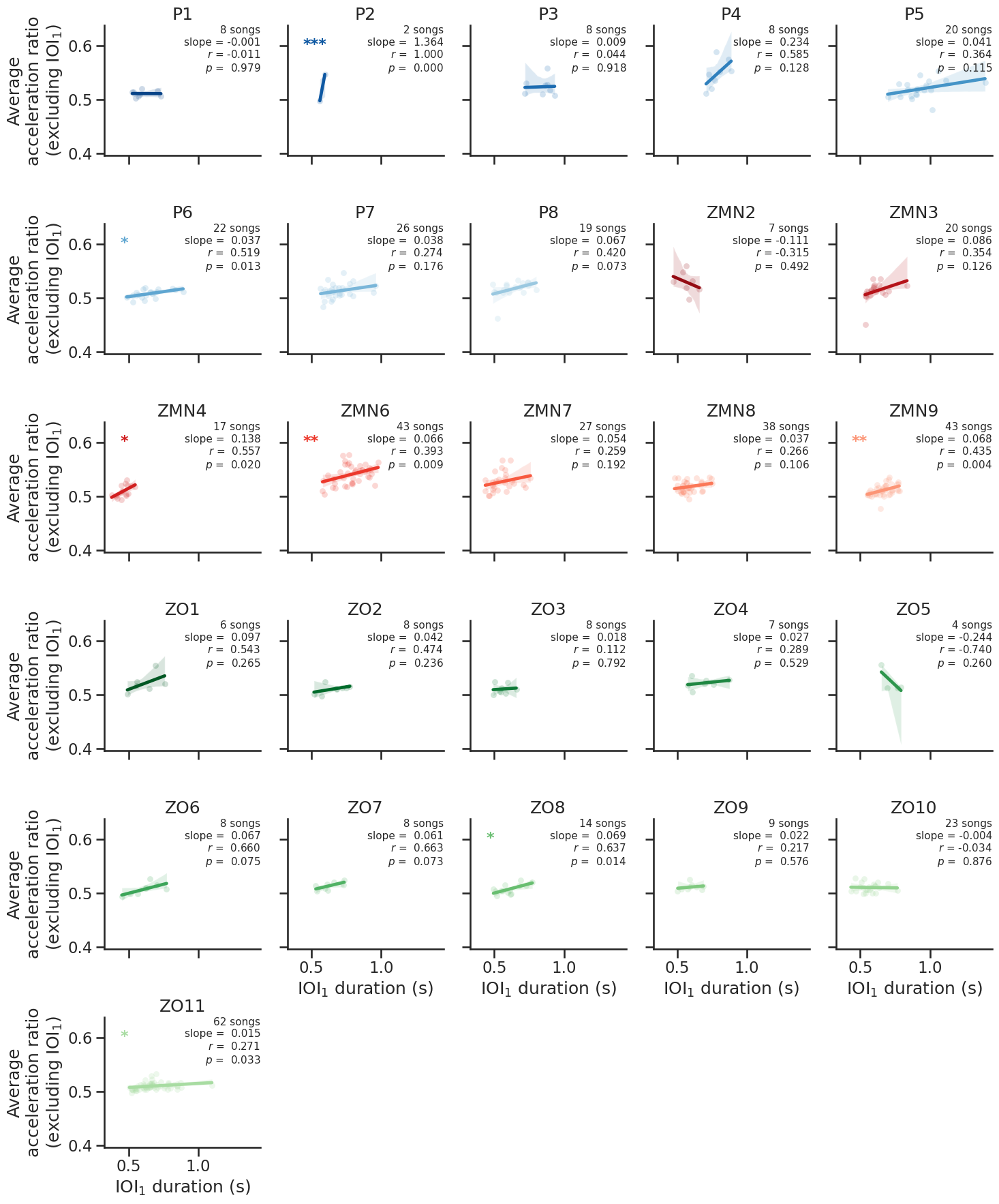
*Figure S8. The top five songs of penguin ZMN6 with the longest (top) and shortest (bottom) IOI_1_.*

Songs starting with a long-duration IOI_1_ tend to have higher acceleration ratios between adjacent IOIs than songs starting with a short-duration IOI_1_. **(A)** The five songs from penguin ZMN6 that have the longest IOI_1_ have **(B)** an average acceleration (after IOI_1_) between approximately 0.52 and 0.56 (horizontal dashed lines). On the contrary, **(C)** the five songs from the same penguin that have the shortest IOI_1_ have **(D)** an average acceleration (after IOI_1_) between 0.50 and 0.54. These exemplars demonstrate the significant effect observed in the main text and Figure 2D: songs with a longer IOI_1_ generally have a higher average acceleration ratio (excluding IOI_1_).

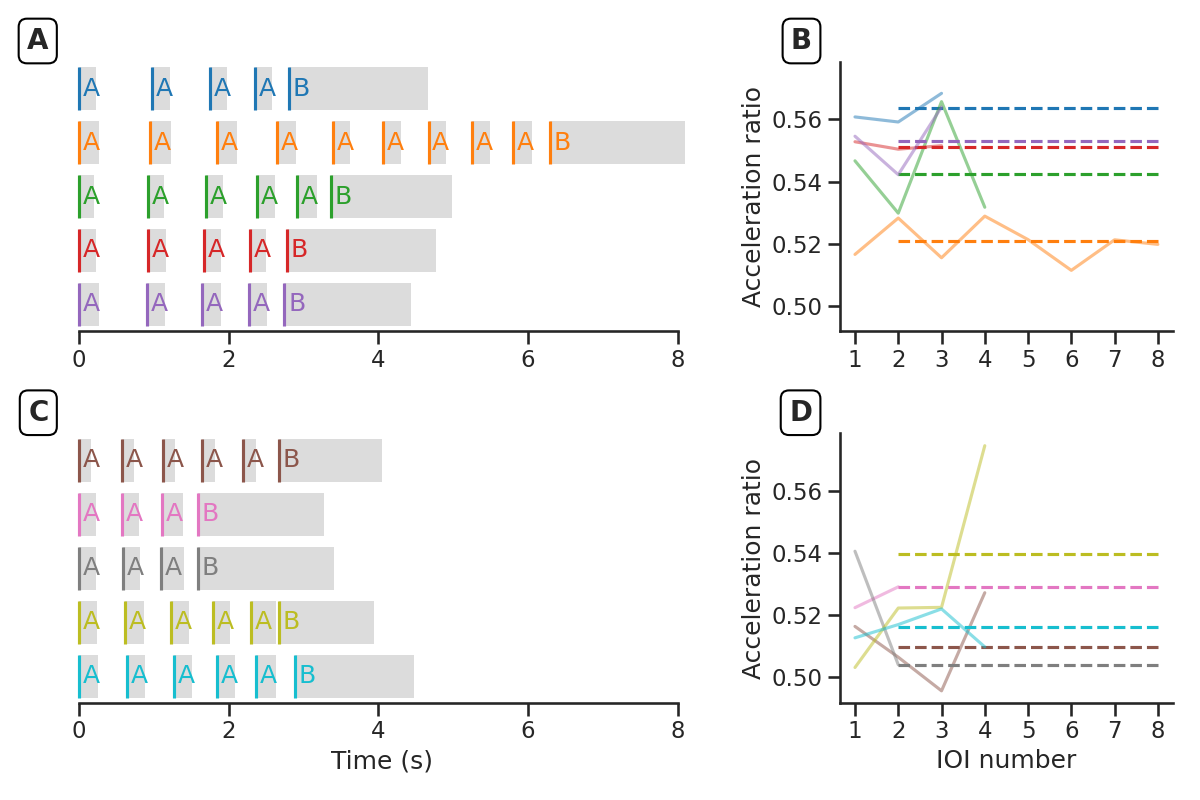

*Figure S9. Per-penguin linear regressions for relative syllable position vs. intensity*

All fitted linear regressions—one per penguin—show a highly significant correlation between relative syllable position and normalized acoustic intensity. This confirms the significant result we find when fitting a single LMEM on the combined data (see main manuscript). Colors (Figure 2) and alphanumeric codes (Table S1) denote penguin identity.

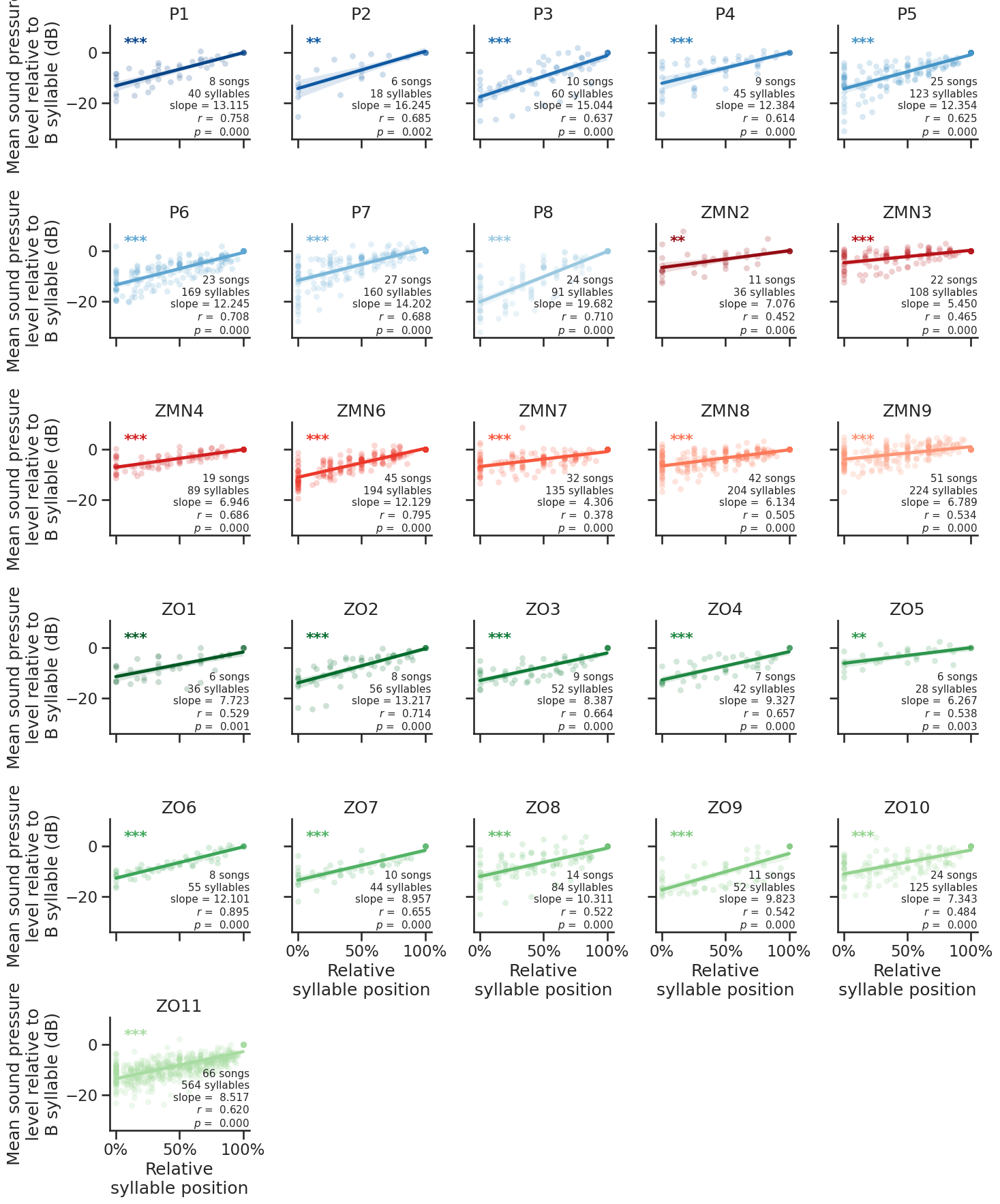

*Supplementary references*

1.	Favaro L., L. Ozella & D. Pessani. 2014. The vocal repertoire of the African penguin (*Spheniscus demersus*): Structure and function of calls. *PLoS One* **9**: e103460. https://doi.org/10.1371/journal.pone.0103460

2.	Favaro L., M. Gamba, E. Cresta, *et al.* 2020. Do penguins’ vocal sequences conform to linguistic laws? *Biology Letters* **16**: 20190589. https://doi.org/10.1098/rsbl.2019.0589

3.	Baciadonna L., C. Pasquaretta, V. Maraner, *et al.* 2024. Network social dynamics of an ex-situ colony of African penguins following the introduction of unknown conspecifics. *Applied Animal Behaviour Science* **273**: 106232. https://doi.org/10.1016/j.applanim.2024.106232

4.	Figel T., S.P. Coyne & K. Martin. 2023. Sex and age differences in activity budgets in a population of captive African penguins (<i>Spheniscus demersus<i/>). *Journal of Applied Animal Welfare Science* **26**: 438–446. https://doi.org/10.1080/10888705.2021.1984916
